# A cortical surface template for human neuroscience

**DOI:** 10.1101/2023.03.21.533686

**Authors:** Ma Feilong, Guo Jiahui, M. Ida Gobbini, James V. Haxby

**Author notes:** Corresponding authors: Ma Feilong, James V. Haxby.

## Abstract

Neuroimaging data analysis relies on normalization to standard anatomical templates to resolve macroanatomical differences across brains. Existing human cortical surface templates sample locations unevenly because of distortions introduced by inflation of the folded cortex into a standard shape. Here we present the onavg template, which affords uniform sampling of the cortex. We created the onavg template based on openly-available high-quality structural scans of 1,031 brains—25 times more than existing cortical templates. We optimized the vertex locations based on cortical anatomy, achieving an even distribution. We observed consistently higher multivariate pattern classification accuracies and representational geometry inter-subject correlations based on onavg than on other templates, and onavg only needs 3⁄4 as much data to achieve the same performance compared to other templates. The optimized sampling also reduces CPU time across algorithms by 1.3%–22.4% due to less variation in the number of vertices in each searchlight.

## Introduction

Various functions of the cerebral cortex are systematically organized on this highly-folded surface ^1–4^. Functional magnetic resonance imaging (fMRI) data, which were acquired as 3D volumes, can be projected onto this surface for analysis and visualization in a 2D space ^5^. Compared with the 3D volumetric analysis of fMRI data, surface-based analysis affords better inter-subject alignment, higher statistical power, more accurate localization of functional areas, and better brain-based prediction of cognitive and personality traits ^6–13^. Due to these advantages, surface-based analysis has been widely adopted by the neuroimaging community, including software ^14–17^, large-scale datasets ^18–20^, and cortical atlases and parcellations ^21–25^.

To account for individual differences in macroanatomy, it is key to normalize all subjects’ data based on an anatomical template, so that the cortical mesh comprises the same number of vertices across brains, and the same vertex corresponds to the same macroanatomical location. The most commonly used template spaces are fsaverage ^5^ and fs_LR ^26^, which were created based on 40 brains. In these standard spaces, the locations of cortical vertices are not based on the anatomical surface, but rather on the spherical surface—a surface obtained by fully inflating each cortical hemisphere to a sphere (Figure 1f). Then, a geodesic polyhedron—usually a subdivided icosahedron—is used to define the locations of cortical vertices. This procedure allows the vertices to be approximately uniformly distributed on the spherical surface.

**Figure 1.**
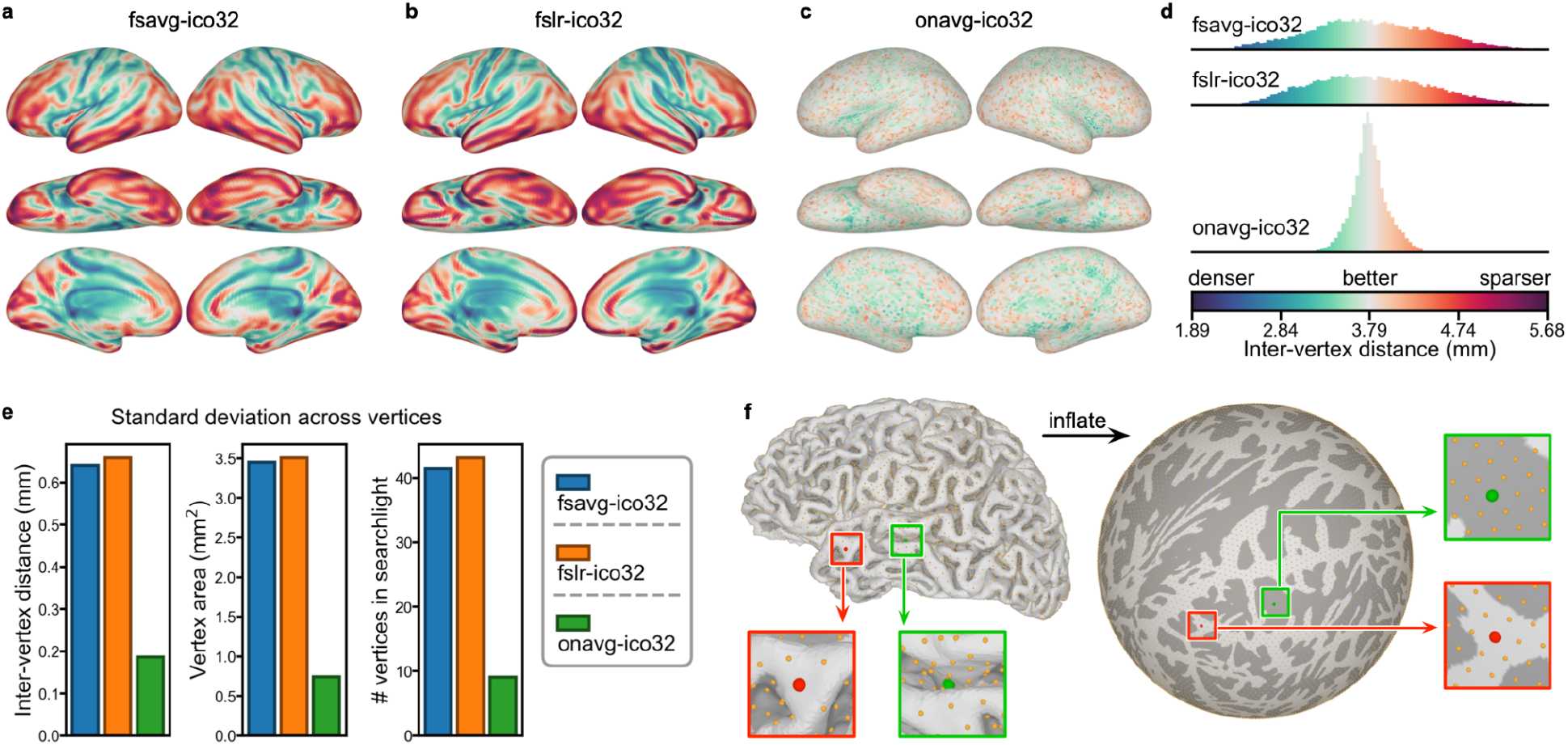
Variation in vertex properties across the cortex. The way classic surface templates sample the cortex is far from uniform. The distribution of vertices is much denser in major sulci compared to other cortical regions for both fsavg (a) and fslr (b), as measured by inter-vertex distance. The new onavg template (c) aims to uniformly sample the cortical surface, and its variation of inter-vertex distance is much smaller compared to previous templates (d). (e) Besides inter-vertex distance, the onavg template also affords less variation in other vertex properties across the cortex, including both the cortical area occupied by a vertex and the number of vertices covered by a 20 mm searchlight. (f) Classic surface templates sample the cortical surface based on the spherical surface, which was obtained by fully inflating the original anatomical surface. For these templates, the distribution of vertices is almost uniform on the spherical surface (right), but far from uniform on the anatomical surface (left), due to the geometric distortion introduced by inflation. Vertices of the same color (red/green; also in zoomed-in views) are homologous for the two surfaces.

However, because the geometry of the spherical surface differs from the original surface, the distribution of cortical vertices is far from uniform on the original anatomical surface. For example, cortical vertices are much denser in the central sulcus and the lateral sulcus than in ventral temporal and prefrontal cortices (Figure 1a, 1b).

In this work, we present the onavg (short for OpenNeuro Average) surface template, a human cortical surface template that affords uniform sampling of the cortex. The onavg template was created using high-quality MRI scans of 1,031 participants from 30 OpenNeuro datasets ^27^—25 times more participants than previous surface templates ^5, 26^. We optimized the vertex locations of the onavg template based on the cortical anatomy of the 1,031 participants, so that these vertices were evenly distributed on the anatomical surface instead of on the spherical surface, affording uniform sampling of the cerebral cortex.

In a series of analyses based on an independent naturalistic movie-viewing dataset ^28^, we demonstrate the advantages onavg offer to various multivariate pattern analysis (MVPA) techniques ^29–31^. On one hand, the anatomy-based sampling of onavg affords better access to the information encoded in spatial response patterns, leading to higher accuracy for multivariate pattern classification ^32^ and higher inter-subject correlation of representational geometry ^33, 34^. By switching to onavg, the same classification accuracy and inter-subject correlation can be achieved with 3⁄4 of the original number of participants (Figure 2). On the other hand, anatomy-based sampling eliminates extremely large searchlights caused by geometric distortions, leading to consistently reduced computational time for computational algorithms ^30, 35, 36^ that rely on searchlight analysis ^11, 37^. We replicated these analyses using different spatial resolutions, different alignment methods, and different numbers of participants, and we observed consistent results across all conditions (Supplementary Figures S1–S7).

**Figure 2.**
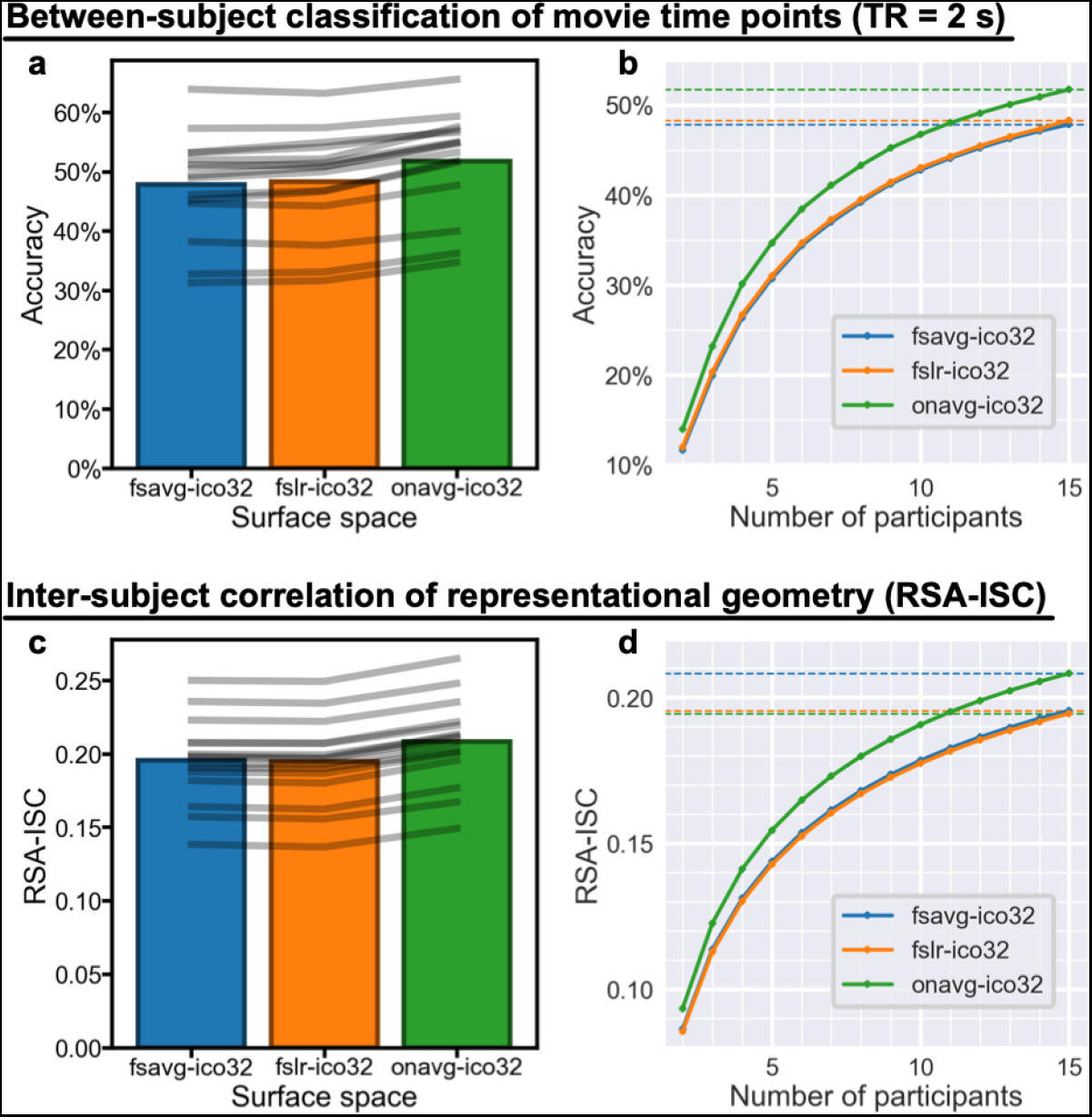
Better cortical sampling improves MVPA results. (a) The between-subject classification accuracy of movie time points based on the onavg surface template (51.7% on average) was consistently higher than the accuracy based on fsavg (47.8%) and fslr (48.3%) across all 15 participants. Gray lines denote individual participants. (b) The accuracy based on 15 participants using fsavg or fslr is approximately the same as the accuracy based on 11 participants using the onavg template (48.0%). Dashed horizontal lines denote accuracies when *n* = 15. (c) RSA-ISC, computed as the correlation between one participant’s RDM and the average of others’, was also higher when using the onavg template (*r* = 0.208, averaged across searchlights) compared with fsavg (*r* = 0.196) and fslr (*r* = 0.195), and the effect was observed for all 15 individual participants. (d) The RSA-ISC based on *n* = 11 and the onavg template (*r* = 0.196) is approximately the same as that based on *n* = 15 and other templates. Across both classification and RSA analyses, we found that the advanced cortical sampling of the onavg template led to more efficient use of fMRI data and better MVPA results, and by switching to onavg the same results can be achieved with less than 3⁄4 of the original sample size. The effect is similar across different sample sizes (Supplementary Figure S7), different resolutions (Supplementary Figure S2), and different alignment methods (Supplementary Figures S4, S5).

## Results

### Not all vertices were created equal

The sphere-based sampling procedure used by traditional surface templates unavoidably leads to inhomogeneous sampling of the cortical surface, as a result of the geometric distortion when each hemisphere was fully inflated to a sphere. The vertices are approximately uniformly distributed on the spherical surface, which means that cortical regions that were expanded during the inflation will be sampled more densely, and cortical regions that were shrunk during the inflation will be sampled more sparsely.

To quantify the distribution of vertices on the cortical surface, we computed the inter-vertex distance for each cortical vertex, where smaller inter-vertex distance indicates denser sampling in the region, and larger distance indicates sparser sampling. For each vertex, we computed the Dijkstra distance between the vertex and its neighbors for each of the 1,031 participants and averaged across participants and neighbors. We performed all our analyses using both the ico32 (also known as icoorder5 or 10k, with mean inter-vertex spacing of approximately 4 mm) and ico64 (icoorder6 or 41k, approximately 2 mm) resolutions and observed consistent results. We focus on the ico32 results in the main text and provide the ico64 results in the supplementary materials.

For both fsaverage and fs_LR (fsavg and fslr for short, respectively, here and thereafter), the inter-vertex distance varied substantially throughout the cortex, and the pattern was similar for both templates (Figure 1a, 1b). The inter-vertex distance was smaller (i.e., denser sampling) in the central sulcus, the postcentral sulcus, the superior temporal sulcus, the lateral sulcus, and much of the cingulate cortex and the medial wall; the inter-vertex distance was larger (i.e., sparser sampling) in the lateral and medial occipital cortex, the lateral and ventral temporal cortex, and the lateral and medial prefrontal cortex (Supplementary Figure S9). In other words, many brain regions that respond in synchronization across subjects ^30, 36, 38–41^ and regions that involve high-level cognition ^42–46^ are not sufficiently sampled based on these traditional surface template spaces.

To resolve these issues caused by traditional templates and sphere-based sampling, we created the onavg surface template using anatomy-based sampling. That is, instead of placing the vertices on the spherical surface based on a geodesic polyhedron, we chose the locations of the vertices based on cortical anatomy: We placed all the vertices on the anatomical surfaces of the 1,031 participants and penalized a pair of vertices if they were too close (see Methods for details). After minimizing the distance-based loss function, the vertices were approximately uniformly distributed throughout the cortex.

The anatomy-based sampling of the onavg template greatly reduced the heterogeneity of vertices in many ways. For inter-vertex distance, the variance across vertices for onavg was 8.41% compared to fsavg, and 7.97% compared to fslr. For the cortical area occupied by each vertex, the variance for onavg was 4.63% compared to fsavg, and 4.49% compared to fslr. For the number of vertices in a 20 mm searchlight, the variance for onavg was 4.73% compared to fsavg, and 4.39% compared to fslr. For all these three vertex properties that we assessed, the variance across the cortex for onavg was much smaller compared to other templates (mean = 5.77%; all < 8.42%).

### Better cortical sampling improves MVPA results

Multivariate pattern analysis (MVPA) comprises algorithms commonly used in computational neuroscience, such as multivariate pattern classification (MVPC) ^29, 32^ and representational similarity analysis (RSA) ^31, 33^. MVPA relies on the fact that the spatial response pattern for a certain stimulus or condition is stable across repetitions within the same subject, or across subjects when their data are functionally hyperaligned ^30, 35, 41^. Therefore, the quality of the spatial patterns formed by cortical vertices is key to successful MVPA.

When resampling neuroimaging data using a traditional sphere-based template, the uneven distribution of vertices on cortical surface creates a systematic bias: Brain regions that have smaller inter-vertex distance are densely sampled and overrepresented, and brain regions that have larger inter-vertex distance are sparsely sampled and underrepresented. Note that oversampling a brain region does not provide extra information—the extra vertices are simply extra interpolations from the same acquired data. In contrast, undersampling does permanently discard certain information, especially the information encoded in fine-grained spatial patterns.

Moreover, each vertex has the same weight when computing the pattern vector, and thus the oversampled regions have more influence on the pattern vector compared to the undersampled regions. In other words, the uneven sampling applies an artificial reweighting to cortical regions based on sampling density, which can affect subsequent computational algorithms.

To assess the effects of different surface templates on MVPA, we performed MVPC and RSA on a naturalistic fMRI dataset ^28^ for each surface space and compared the results. The dataset was collected from 15 participants when they watched the audio-visual movie *Forrest Gump* in a 3 T MRI scanner. We preprocessed the dataset with fMRIPrep ^14^ and functionally hyperaligned all participants’ data using warp hyperalignment ^35^ trained on the first half of the movie. The analysis was performed using the second half of the movie, and therefore the test data was independent of the training data. For similar results using other alignment methods, see Supplementary Figures S4–S7.

In the MVPC analysis, we tried to classify which time point of the movie the participant was watching among all 1,781 time points (TRs; 2 seconds each) based on the brain response patterns. We used a leave-one-subject-out cross-validation and left out a test participant each time. For each time point of the movie, we computed the average response pattern across all other participants as the predicted response pattern of the test participant. Therefore, for each test participant, we had 1,781 measured response patterns and 1,781 predicted response patterns. We examined whether the measured response pattern for a certain time point had the highest correlation to the predicted pattern for the same time point among all 1,781 predicted response patterns (chance accuracy < 0.1%). The classification accuracy based on the onavg template was higher than the accuracies based on fsavg and fslr for all 15 participants, and the average accuracy across all participants was 51.7% (onavg), 47.8% (fsavg), and 48.3% (fslr), respectively (Figure 2a).

The classification accuracy for between-subject MVPC depends on the number of participants. Averaging across a larger number of participants reduces the noise in the predicted response patterns of the test participant, which improves classification accuracy. For all three surface templates, MVPC accuracy consistently increased with more participants (Figure 2b). Note that the same accuracy for fsavg and fslr with *n* = 15 (47.8% and 48.3%, respectively) is approximately the same as the accuracy for onavg with *n* = 11 (48.0%), or *n* = 10.9 and *n* = 11.3, respectively, based on spline interpolation. In other words, the onavg surface template only requires 72.4% and 75.2% of the number of participants for fsavg and fslr, respectively, to achieve the same classification accuracy.

In the RSA analysis, for each searchlight (20 mm), we computed a time-points-by-time-points representational dissimilarity matrix (RDM) for each participant using correlation distance. We computed the Pearson correlation between each participant’s RDM and the average of others’, which is the inter-subject correlation (ISC) of representational geometry ^36^, which is often used as the lower-bound of noise ceiling estimation ^34^. We refer to this correlation as RSA-ISC here and thereafter. We averaged the RSA-ISCs across all searchlights and obtained an average RSA-ISC for each participant. The average RSA-ISC based on the onavg template was consistently higher than the average RSA-ISCs based on fsavg and fslr for all 15 participants, and the average RSA-ISC across all participants was 0.208 (onavg), 0.196 (fsavg), and 0.195 (fslr), respectively (Figure 2c).

Similar to between-subject MVPC accuracy, the RSA-ISC also benefits from the reduction in noise by averaging over a larger number of participants. For all three surface templates, the RSA-ISC consistently increases with more participants (Figure 2d). The same RSA-ISC for fsavg and fslr with *n* = 15 (0.196 and 0.195, respectively) is approximately the same as the RSA-ISC for onavg with *n* = 11 (0.195), or *n* = 11.1 and *n* = 10.9, respectively, based on Spearman–Brown interpolation. In other words, the onavg surface template only requires 74.1% and 72.5% of the number of participants for fsavg and fslr, respectively, to achieve the same RSA-ISC.

In both between-subject MVPC and RSA-ISC, improvements in performance were unevenly distributed, with greater improvement in sparsely-sampled and inhomogeneously-sampled cortical fields (e.g., medial occipital, ventral temporal, premotor, and insular cortices; Figure S8), indicating that other templates bias the anatomical distribution of results from multivariate pattern analyses.

The improvement of MVPC accuracy and RSA-ISC was consistent across individuals—onavg outperformed fsavg and fslr for all 15 participants (Figure 2a, 2c). This was likely because anatomy-based sampling improved every participant’s data. We repeated our analysis using different resolutions (Supplementary Figure S2), different alignment methods (Supplementary Figures S4, S5), and different sample sizes (Supplementary Figure S7), and we observed highly consistent results.

### Expedited computations for searchlight-based algorithms

Searchlight analysis ^11, 37^ is widely used in combination with MVPC or RSA to assess which part of the brain contains the information of interest, and it serves as the backbone of computational algorithms such as searchlight hyperalignment ^35, 36, 38^. Each time, a brain region (i.e., a searchlight) is chosen to perform the analysis in, and the region is defined as the group of vertices that are within a certain distance (i.e., searchlight radius) from a given searchlight center. Traditional surface templates have large variation in inter-vertex-distance across the cortex, and, as a result, large variation in the number of vertices in a searchlight (Figure 1e). The densely sampled brain regions have more vertices in each searchlight, causing prolonged computations in these searchlights.

We systematically assessed the computational time for various searchlight-based algorithms, and we observed a consistent effect that the computational time based on the onavg surface template was shorter than the computational time based on fsavg and fslr, with a 1.3%–24.4% reduction in CPU time (Figure 3). For creating the common space for hyperalignment—an “average” participant with representative representational geometry and functional topography—the CPU time was 88.5% and 89.4% compared with fsavg and fslr, respectively. For hyperalignment of each participant to the common space, the CPU time was 75.6% and 79.3% compared to fsavg and fslr, respectively, based on the classic Procrustes algorithm ^41^, and 83.9% and 85.9%, respectively, based on the warp hyperalignment algorithm ^35^. For searchlight-based RSA analysis, the CPU time was 92.8% and 94.7% compared to fsavg and fslr, respectively. For searchlight-based classification analysis, the CPU time was 96.7% and 98.7% compared to fsavg and fslr, respectively. On average across conditions, switching to onavg led to a 11.5% reduction in CPU time.

**Figure 3.**
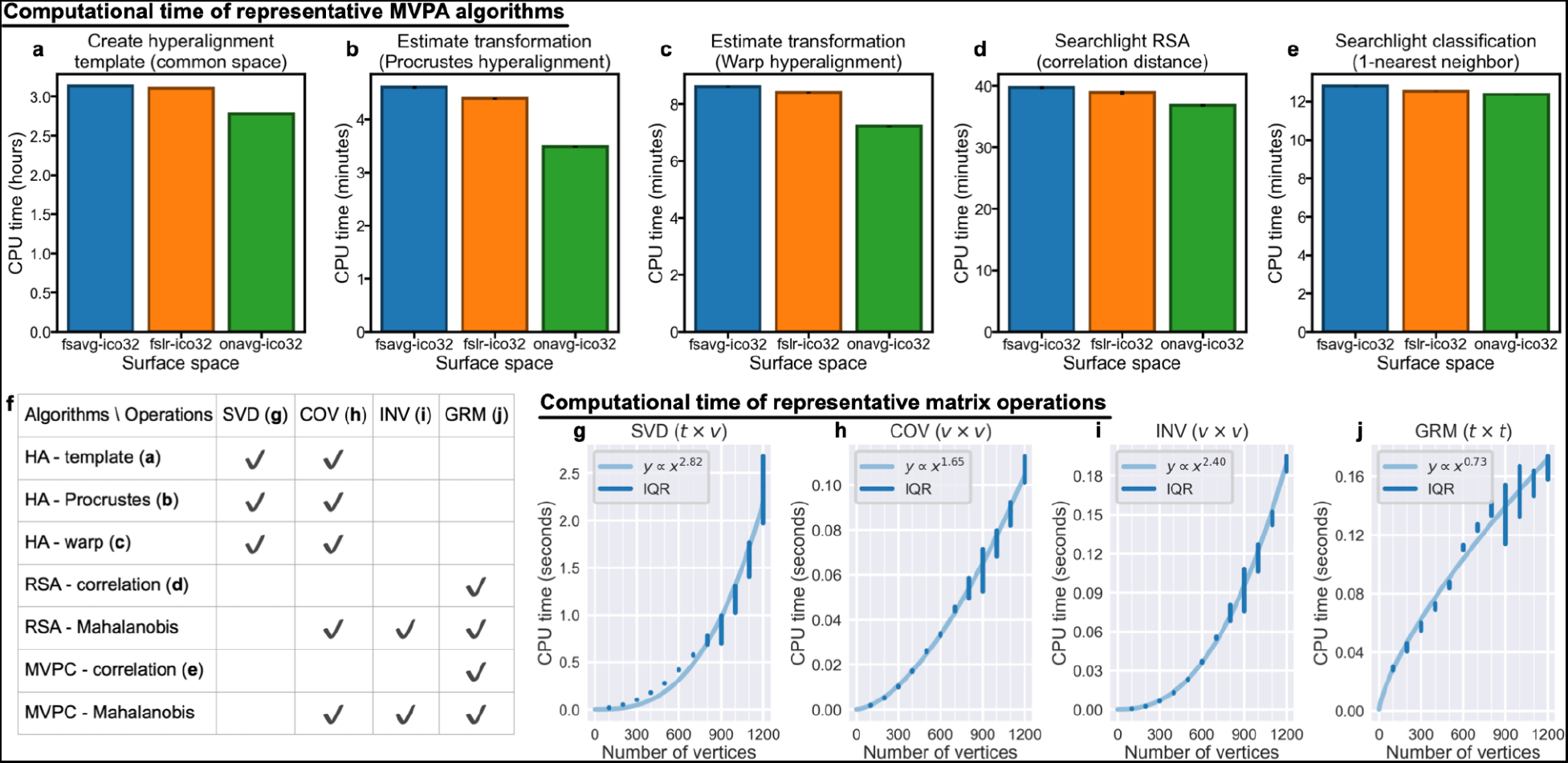
Computational time of representative computational algorithms. The anatomy-based uniform sampling of the onavg template avoids sizable searchlights (Figure 1e), resulting in expedited computations for various searchlight-based algorithms, including creating hyperalignment common space (a), aligning individual subjects to the common space (b and c), representational similarity analysis (d), and multivariate pattern classification (e). This is because these algorithms rely on basic matrix operations (f), and the computational time of matrix operations grows exponentially with searchlight size (g–j). The computational time is approximately proportional to a power of the number of vertices in the searchlight (light blue curves). The exponent varies between 0.73 and 2.82 for different matrix operations, which means doubling the number of vertices will make the computations 7.07, 3.14, 5.29, and 1.66 times as long, for SVD, COV, INV, and GRM, respectively. The vertical blue lines denote the interquartile range (IQR) of 100,000 repetitions. SVD, singular value decomposition; COV, covariance matrix; INV, inverse of covariance matrix; GRM, Gram matrix. In the top row, black error bars denote standard error of the mean, which are sometimes too small to be visible.

The reduction in CPU time is because these computational algorithms rely on matrix operations of the input data matrix (Figure 3f), and the time required by these matrix operations grows exponentially with the number of vertices (Figure 3g–j). As a result, a large searchlight will lead to prolonged matrix operations, and eventually, prolonged CPU time for computational algorithms. For example, if the computational time is proportional to the number of vertices squared (i.e., quadratic complexity), doubling the number of vertices requires 4 times as much computational time. For representative matrix operations, the exponent varies between 0.73 and 2.82, and the computations are up to 7.07 times as long when the number of vertices is doubled (Figure 3g–j). The onavg template avoids sizable searchlights created by geometric distortions and uneven sampling (Figure 1e), and therefore it avoids the unnecessary prolonged computations and speeds up computational algorithms substantially.

## Discussion

In this work we introduce a new cortical surface template, “onavg”, that was built to achieve uniform sampling of cortical vertices based on high-quality structural scans of 1,031 brains. Compared to classic templates that rely on sphere-based sampling, onavg greatly facilitates the efficient use of neuroimaging data and improves the results of various MVPA algorithms. Furthermore, onavg avoids sizable searchlights caused by uneven sampling and expedites computational methods based on searchlight analysis.

Replicability and reproducibility are key to neuroscientific research ^47–51^, and one of the best practices to increase replicability and reproducibility is to use larger amounts of data ^52–54^. However, this is often infeasible in practice due to the cost and human effort needed to collect and curate fMRI data. Alternatively, making more efficient use of existing data also increases statistical power, and in turn, better replicability and reproducibility ^44, 55^. The onavg template provides a new way to make better use of neuroimaging data in surface-based analysis. It consistently improved the results of MVPA algorithms across different subjects, different data resolutions, different alignment methods, and different amounts of data (Figure 2, Supplementary Figures S2, S4–S7). Compared with commonly used surface templates, onavg only requires 3/4 of the amount of data to achieve the same level of performance. Therefore, onavg has a great potential to both improve the replicability and reproducibility of future neuroscientific research and reduce the cost and effort of neuroimaging data collection.

Besides improved performance, onavg also reduces the computational time of various MVPA algorithms (Figure 3, Supplementary Figure S3), which highly depend on the number of vertices in a searchlight. The geometric distortion of sphere-based sampling creates large searchlights in densely sampled regions, which can be avoided by switching to anatomy-based sampling, which we used to create onavg (Figure 1e).

Therefore, onavg avoids prolonged computations in large searchlights. Due to the size of movie time point RDMs (1,781 × 1,781), we only benchmarked the RSA computational time based on correlation distance, whose effect size might be smaller than those of alternative distance metrics. For example, the crossnobis distance ^56^ requires the computation of the covariance matrix and its inverse (Figure 3f). These operations have higher time complexity, which could benefit more from avoiding the sizable searchlights created by geometric distortion.

The onavg template was created based on high-quality structural scans of 1,031 brains, more than 25 times more brains as compared to previous surface templates. This was a direct benefit from open science, especially the datasets hosted on OpenNeuro ^27^ and managed by DataLad ^57^. We have made the onavg template openly available under the Creative Commons CC0 license and released it as a DataLad ^57^ Dataset on both OSF (https://osf.io/uprh3/) and GitHub (https://github.com/feilong/tpl-onavg). The onavg template is also being integrated into TemplateFlow ^58^, the standard repository for neuroimaging templates.

## Methods

### Datasets

We built the onavg surface template based on high-quality structural scans of 1,031 brains aggregated from 30 OpenNeuro datasets ^27^. The datasets and participants were selected based on a few criteria:

1. The participant has at least a high-quality T1-weighted scan and a high-quality T2-weighted scan. That is, the T1-weighted and T2-weighted scans both have (a) whole coverage of the cerebral cortex, (b) a spatial resolution of 1 mm or less in all directions, and (c) no major quality issues.
2. The participant’s structural scans show no visible lesion or abnormality. For ‘ds002799’ only pre-op scans are included.
3. The structural workflow of fMRIPrep successfully finishes within 100 hours of CPU time and without errors and the reconstructed cortical surface has no major artifacts. Longer sessions are predominantly caused by prolonged ‘mris_fix_topology’, and that usually indicates problematic structural images.
4. The dataset is released under CC0 or PDDL license.

After screening, 1,154 participants passed our criteria. We noted that some of the participants were duplicates. For example, the same individual might have participated in multiple experiments and appear in multiple datasets. To find out these duplicates, we compared the similarities of reconstructed cortical surfaces and found 1,031 unique participants. The duplicate participants identified by reconstructed surfaces are consistent with the documentation of the corresponding datasets.

The duplicates can be summarized into two main categories:

1. The same participant participated in multiple conditions/sessions of the same dataset, and these different conditions/sessions from the same participant were coded using different subject IDs. For example, in ‘ds003242’, ‘sub-SAXSISO01b’, ‘sub-SAXSISO01f’, and ‘sub-SAXSISO01s’ are the same participant; in ‘ds003849’, ‘sub-BPD0050À and ‘sub-BPD0050B’ are the same participant; in ‘ds001597’, ‘sub-07’ and ‘sub-17’ are the same participant; in ‘ds002320’, ‘sub-51’ and ‘sub-88’ are the same participant.
2. The same participant appeared in different datasets. For example, ‘sub-f1031ax’ in ‘ds003452’ and ‘sub-f1031ax’ in ‘ds003465’ are the same participant; ‘sub-09’ in ‘ds001597’ and ‘sub-14’ in ‘ds001233’ are the same participant; ‘sub-MSC06’ in ‘ds000224’ and ‘sub-cast2’ in ‘ds002766’ are the same participant. For ‘ds003452’ and ‘ds003465’, the same participant also has the same subject ID.

During analysis, we averaged the multiple reconstructed surfaces of the same participant and created an average surface for each participant.

### Preprocessing and surface reconstruction

We downloaded and managed the data files of the 30 OpenNeuro datasets using DataLad ^57^, and preprocessed the structural scans of these 1,031 participants using fMRIPrep ^14^ version 21.0.1. Specifically, we used the ‘--anat-only’ option and designated fsaverage as the output space. These settings allowed us to reconstruct the cortical surfaces of these participants using FreeSurfer ^15^ while benefiting from the optimized structural preprocessing workflow of fMRIPrep. To increase the replicability of our results, we also used the ‘--skull-strip-fixed-seed’ option with a random seed of 0. This ensures that anyone could regenerate identical cortical surfaces as those used in this work.

FreeSurfer also generated automatic segmentation of the brain (“aseg” files) ^59^, which we used to verify that the brain was properly covered by the structural images. Specifically, if the cortical gray matter appeared around the boundary of the structural image based on the “aseg” parcellation, we examined the structural image again manually to make sure this was not because of partial coverage of the cortex.

### Optimize the template using anatomy-based sampling

We optimized the vertex locations of the onavg template, so that no vertices were too close to each other, and the vertices were approximately uniformly distributed throughout the cortex. In the optimization, we used a distance-based objective function, which penalizes pairs of vertices if they were too close. We first performed a coarse discrete optimization based on a geodesic grid, which chose a set of vertex locations that minimized the loss function from a larger set of candidate locations. We then performed a fine optimization, which allowed the vertices to move freely nearby in small steps to further reduce the loss function.

#### Objective function

We defined the objective function (loss function) of the optimization process as:

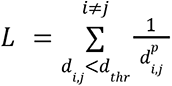

where *d_i,j_* is the Dijkstra geodesic distance between a pair of vertices *i* and *j*. The distance was based on the anatomical surfaces of all 1,031 participants, and therefore the objective of the optimization process was to make the vertices evenly distributed on the anatomical surface instead of the spherical surface. This distance was raised to a power of *p* to further penalize small distances (i.e., neighboring vertices being too close). In practice we used *p* = 4, which worked well, thus we didn’t explore other options. Our objective was to ensure the distance was not too small for vertices close to each other, and it is computationally heavy to compute and manage all pairwise distances between vertices. Therefore, we employed a cutoff distance *d_thr_* and only included vertex pairs whose distance was smaller than the cutoff distance *d_thr_*. We chose *d_thr_* to be 256 mm / ico, which was 8 mm for ico32, and 4 mm for ico64, approximately twice the average inter-vertex distance.

Occasionally, two vertices might appear at the same location during the optimization process, which makes the distance zero and the loss function ill-defined. To avoid this problem, we added a small number *ε* to the distance *d_i,j_* in the steps where this problem might happen, and we used *ε* = 0.001 in practice.

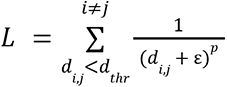

#### Coarse optimization of vertex locations

The coarse optimization was a discrete optimization, where we chose the vertex locations from a large set of candidate locations, so that the loss function was minimized. The candidate locations were the vertex locations of a high-resolution reference surface. We created the high-resolution reference surface by upsampling the fsaverage spherical surface to a higher resolution. Specifically, we used fsavg-ico256 (655,362 vertices per hemisphere) to optimize onavg-ico32 (10,242 vertices per hemisphere), and fsavg-ico512 (2,621,442 vertices per hemisphere) to optimize onavg-ico64 (40,962 vertices per hemisphere). In other words, the locations of the 10,242 vertices were chosen from the 655,362 candidate locations, and the locations of the 40,962 vertices were chosen from the 2,621,442 candidate locations. The number of candidate locations was approximately 64 times the number of vertices.

We initialized the vertex locations by randomly choosing candidate locations without replacement. Then, we tried to find better locations for them. Each candidate location had a loss value based on which vertex locations near it had been occupied, and this value was the same value that would be added to the loss function if the location was occupied by a vertex. For each vertex, we first removed it from its current location and updated the loss value of all candidate locations. We then placed the vertex to the location that had the minimal loss value. We looped through all vertices for up to 100 times and updated their locations accordingly. This process might have stopped early if the local optimum was reached before 100 iterations. The order of vertices was randomized during each iteration.

This coarse optimization process was a greedy algorithm. The local minimum might not be the global minimum, and the results depended on initialization. Therefore, for each hemisphere and each resolution, we repeated the process for 200 times with different random seeds (and different initializations accordingly). We chose the one that had the smallest loss value for further optimization.

#### Fine optimization of vertex locations

We refined the vertex locations after the coarse optimization to further reduce the loss function. This time, instead of predefined locations, we allowed the vertices to move freely nearby. For each vertex, we used numerical differentiation to find the direction of the gradient, and we moved the vertex along the direction to reduce the loss function. We computed new loss values for different step sizes ranging from 2^-21^ to 2^-10^ (4×10^-7^ and 1×10^-3^) and used the optimal step size multiplied by 0.5 as the final step size to update the vertex location. The factor of 0.5 was because the optimization was performed simultaneously across vertices in parallel, and if the optimal was to reduce the distance between two vertices by 1 mm, each of them should only be moved by 0.5 mm.

It is difficult to compute the Dijkstra distance in this case, because the vertex locations of the new surface don’t correspond to vertex locations of the high-resolution reference sphere. Therefore, we approximate the distance based on barycentric interpolation. Each vertex is located on a face of the triangular mesh, and its coordinates ***c****_i_* can be represented as a weighted sum of the three vertices of the triangle.

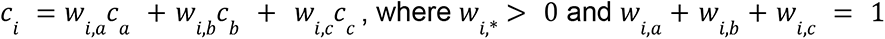

Similarly, say vertex *j* locates on a triangle whose vertices were *x*, *y*, and *z*:

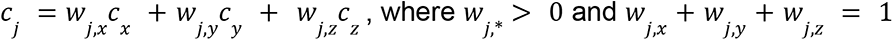

We estimate *d_i,j_* as

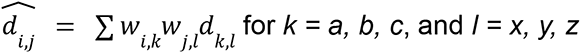

This allowed us to compute the distance between a pair of vertices at any locations and further fine-tune the vertex locations without being constrained by the reference sphere.

#### Optimization of triangular faces

After finalizing the vertex locations, we created an initial surface mesh based on these vertices. Specifically, we created a convex hull based on the vertex locations on the spherical surface, and the simplices of the convex hull were the triangular faces of the initial surface mesh.

We wanted to make each triangular face as similar to an equilateral triangle as possible, and therefore we optimized the faces to avoid long edges and elongated triangles. Each pair of neighboring faces forms a quadrilateral *ABCD*. When *AC* < *BD*, we divide the quadrilateral into two triangles *ABC* and *ACD*; when *AC* > *BD*, we divide the quadrilateral into two triangles *ABD* and *CBD*. Note that the edge lengths were computed based on the anatomical surface of the 1,031 participants, rather than the spherical surface. We repeated this procedure until no further optimization can be performed.

For each triangular face, we also changed the order of its three vertices, *A*, *B*, and *C*, so that the cross product of *AB* and *BC* is the same direction as the outward normal of the face. The purpose of the step was to make it easier to compute surface normals and make the generated faces more compatible with those generated by FreeSurfer (https://surfer.nmr.mgh.harvard.edu/fswiki/FreeSurferWiki/SurfaceNormal).

Note that the optimization of triangular faces does not affect the vertex locations, the interpolated data, or the MVPA results. The purpose of the optimization was simply to make the triangular faces of the surface mesh better describe the geometry of the cortical surface.

### Template evaluation

#### Inter-vertex distance and other vertex properties

The cortical surface mesh comprises a set of cortical vertices, and the vertices are connected by edges, forming triangular faces. For each vertex, we define its neighbors as the vertices connected to it by an edge. We computed the distance between each vertex and its neighbors and averaged across all 1,031 participants and all neighbors. We used this average distance as the inter-vertex distance of the vertex. Therefore, the inter-vertex distance measures the density of vertices in a local area, where smaller inter-vertex distance indicates denser vertices, and larger inter-vertex distance indicates sparser vertices. To evenly sample the cortex, the inter-vertex distance should have minimal variation across all vertices.

We computed the area of each triangular face and divided it equally among the three vertices of the face. In other words, the area occupied by each vertex was a third of the area of all faces comprising the vertex. Therefore, smaller vertex area indicates denser vertices, and larger vertex area indicates sparser vertices. Similar to inter-vertex distance, ideally the variation of vertex area should be as small as possible.

For each vertex, we created a searchlight around it, which was the group of vertices that had a geodesic distance of 20 mm or less from the center vertex. The geodesic distance was computed as the average of all 1,031 participants. The number of vertices in a searchlight varies by brain region—the number is larger for regions with denser vertices, and smaller for regions with sparser vertices.

All these three vertex properties (inter-vertex distance, vertex area, and number of vertices in a 20 mm searchlight) measures the density of vertices in a local brain area. We expect these properties to have larger variation when the cortex is sampled unevenly, and smaller variation when the cortex is sampled evenly. We computed the standard deviation of these properties and compared them across different surface templates, and we found the onavg template had much smaller standard deviation compared to other templates (Figure 1).

#### Test dataset for MVPA algorithms

We used the *Forrest* dataset ^28^ to evaluate the surface templates. The dataset was part of the phase 2 data of the studyforrest project (https://www.studyforrest.org/), and it includes fMRI data of 15 participants that were collected while the participants watched the feature movie *Forrest Gump*. We preprocessed the dataset with fMRIPrep ^14^ and resampled them to different surface spaces. The movie was approximately 2 hours long, and during the scan it was divided into 8 runs. We used the first half of the movie (the first 4 runs; 1,818 TRs in total; TR = 2 seconds) to train hyperalignment models, and the second half of the movie (1,781 TRs) to perform the main analysis. Note that the *Forrest* dataset was not among the 30 OpenNeuro datasets that we used to create the template, and therefore it’s completely independent of the template creation process.

#### Hyperalignment template creation

For each surface template space, we created a hyperalignment template, so that all subjects’ data can be projected into this common template space. In the common template space, idiosyncrasies in functional–anatomical correspondence are resolved, and response patterns can be compared across participants (Figure 2; Supplementary Figures S4 and S5). We followed the procedure described in Feilong et al. (2022) ^35^ to create the template. We first created a local template for each searchlight (20 mm radius), and we made both the representational geometry and the topography of the local template reflective of the central tendency of the group of participants. We then aggregated the local templates and formed a template for each hemisphere, and we combined both hemispheres’ templates into a whole-cortex template. This template creation process made heavy use of principal component analysis (PCA) and the orthogonal Procrustes algorithm ^41^, which rely on singular value decomposition (SVD) and the computation of covariance matrices (COV).

#### Hyperalignment to template

For each surface template space, we prepared three sets of data based on different alignment methods: surface alignment (i.e., no hyperalignment), Procrustes hyperalignment ^36^, and warp hyperalignment ^44^. We performed all hyperalignment training based on the first half of the movie data and estimated the hyperalignment transformations. We then applied these transformations to the test data (second half of the movie), which was independent of the training data. We report the results based on warp hyperalignment in the main text (Figure 2), and the results based on Procrustes hyperalignment and surface alignment in Supplementary Figures S4 and S5, respectively. The classification accuracy and RSA-ISC were both higher for warp hyperalignment than Procrustes hyperalignment and surface alignment, as a result of better alignment across individuals. The differences between surface templates were similar for all three alignment methods, and onavg consistently outperformed other surface templates.

Both Procrustes hyperalignment and warp hyperalignment used in this study are based on searchlight hyperalignment. For each participant, we obtained a local transformation for each searchlight (20 mm radius) and combined these local transformations to form a whole-cortex transformation. The estimation of the transformation made heavy use of ridge regression and the orthogonal Procrustes algorithm ^41^, which rely on singular value decomposition (SVD) and the computation of covariance matrices (COV).

#### Multivariate pattern classification of movie time points

For each surface template space and each alignment method, we performed a between-subject multivariate pattern classification of movie time points (TRs). We used a leave-one-subject-out cross-validation and a nearest neighbor classifier. We also trained a PCA model based on the first half of the movie (training data) and applied it to the second half of the movie (test data), so that the classification was based on normalized PCs. The number of PCs was also chosen based on the first half of the movie with a nested cross-validation. The test data comprises 1,781 time points, and therefore each participant had 1,781 measured brain response patterns, one for each time point. Each time, we left out a test participant and computed 1,781 predicted response patterns of the test participant, one for each time point, by averaging the response patterns across other participants. For each measured response pattern, we computed its correlation with all 1,781 predicted response patterns and predicted which time point the participant was watching based on which predicted pattern had the highest correlation. In other words, there were 1,781 choices for this classification task, and the classification was only correct if the corresponding predicted pattern had the highest correlation with the measured pattern. Therefore, the chance level was less than 0.1%. In practice, this classification task can be performed using a correlation-based similarity matrix (1,781 × 1,781), which is a Gram matrix based on the normalized response patterns.

A successful classification relies on the quality of the predicted patterns, and the quality can be improved by averaging over a larger amount of data (i.e., averaging over more training participants), which reduces the noise relative to signal. We repeated the classification analysis with smaller numbers of participants, and correspondingly, smaller numbers of training participants. There are multiple ways to choose a subset of participants from the entire set of 15 participants, and therefore, for each number of participants, we repeated the sampling procedure for 100 times with different random seeds, and we averaged the results across the 100 repetitions.

#### Inter-subject correlation of representational geometry

Similar to the classification analysis, we repeated the RSA analysis for each surface template space and each alignment method. The RSA analysis was a searchlight analysis. For each searchlight (20 mm radius), we computed a time-point-by-time-point RDM for each participant based on correlation distance. The RDM was based on the test data (second half of the movie), and it comprised 1,781 × 1,781 elements. We computed the inter-subject correlation of representational geometry as the correlation between one participant’s RDM and the average of others’, which we refer to as RSA-ISC. For each left out test participant, we averaged the RSA-ISC across all searchlights and obtained a single average correlation. When we averaged across multiple correlation coefficients, we used the Fisher-transformation to transform the correlation coefficients to zs, which are approximately normally distributed, and we transformed it back after averaging.

Similar to the classification analysis, the quality of an RDM can be improved by averaging over larger numbers of participants. The quality of the RDM is often measured by its reliability using Cronbach’s alpha coefficient. Furthermore, based on the Spearman–Brown prediction formula, we can estimate how this reliability changes with the number of participants used in averaging.

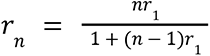

where *r_n_* is the reliability of the RDM obtained by averaging over *n* RDMs.

In this formula, *r_1_* can be estimated using *r_n_* and *n*, and in this case *n* = 14 (15 participants in total, one left out). After obtaining *r_1_*, we can use it to estimate *r_n_*for different choices of *n*. By combining Cronbach’s alpha coefficient and the Spearman–Brown prediction formula, we estimated the reliability of the average RDM, for different numbers of participants.

The correlation between the average RDM and the left out participant’s RDM is proportional to the square root of the average RDM’s reliability. Therefore, by estimating the average RDM’s reliability, we can estimate the correlation between the two RDMs for smaller numbers of participants (Figure 2d).

#### Computational time of MVPA algorithms

For all the MVPA algorithms that we performed, we recorded the CPU time with Python’s ‘time.process_time_ns’ function, which affords nanosecond resolution. In this work, we ran the algorithms in single processes and made sure that the measured CPU time was accurate. In scenarios where recording CPU time is not necessary, it is often better to use parallel computing (e.g., with Python’s ‘joblib’ package), which reduces the walltime of these algorithms significantly. For the same algorithm, the CPU time of different surface templates was computed on the same node of Dartmouth’s Discovery cluster to eliminate potential confounds from hardware and software differences. We repeated each algorithm for different surface template spaces and recorded the CPU time accordingly.

For each surface template space, we created a hyperalignment template for each hemisphere and recorded the CPU time. We summed over the CPU time across both hemispheres and obtained a total CPU time for each surface template space (Figure 3a). When we estimated the hyperalignment transformations, we recorded the CPU time for each participant and each hemisphere. Similar to hyperalignment template creation, we computed the sum of the CPU time across both hemispheres and obtained a total CPU time for each participant. We used two different hyperalignment algorithms in our analysis, and therefore we repeated this process for each algorithm (Figure 3b and 3c, respectively).

We also performed searchlight classification and searchlight RSA for each template space. The searchlight classification analysis was similar to the whole cortex classification analysis, except each time the data was from a 20 mm searchlight instead of PCs of the entire cortex, and we classified 5-TR segments (10 s each) instead of single TRs (2 s each). We recorded the CPU time for each participant and each hemisphere, and added together the CPU time of both hemispheres. For the searchlight RSA analysis, we recorded the total CPU time of all participants for each searchlight. This was because estimating the RDM noise ceiling requires all participants’ RDMs, and it is impractical to save these RDMs, and therefore we performed the analysis and recorded the CPU time for each searchlight separately, which does not require saving RDMs to storage. We performed searchlight classification and searchlight RSA for all three alignment methods and averaged the CPU time. This was because for the same surface template, the data matrix shape and the searchlights were the same across different alignment methods, and thus the theoretical computational complexity was the same. By averaging across these repetitions we further reduce the noise in measured CPU time.

#### Computational time of basic matrix operations

Complex computational algorithms are based on basic matrix operations (Figure 3f). For example, Procrustes hyperalignment relies on singular value decomposition (SVD) of the covariance matrix (COV); correlation-based RSA relies on computing the Gram matrix (GRM); RSA with alternative distance metrics, such as the crossnobis distance, requires the inversion (INV) of the covariance matrix.

The computational time of these matrix operations does not grow linearly with the number of vertices, instead, it takes much longer when the number of vertices is large. To better demonstrate this effect, we systematically evaluated the CPU time of these matrix operations as a function of the number of vertices of the data matrix. We generated random data matrices with different numbers of vertices, ranging from 100 to 1,200 with steps of 100. All these matrices had 1,781 time points, which was the same as the test data. For each number of vertices, we executed these matrix operations 10,000 times each with different random data matrices, which were generated with different random seeds.

To better illustrate the relationship between the CPU time and the number of vertices, we fit an exponential curve *y = x^p^*, where *y* is the CPU time, *x* is the number of vertices, and *p* is the exponent. The exponent *p* is often between 2–3 (Figure 3 g–j).

As a result, if a searchlight contains twice as many vertices compared to the average, the computational time for the searchlight would be 4–8 times as long. When the cortex is unevenly sampled, there are densely sampled regions where the number of vertices is particularly high. Furthermore, there are also more searchlights in these regions, also because the region is densely sampled. As a result, all kinds of searchlight analysis spend prolonged computational time in the densely sampled regions, and when the cortex is evenly sampled, the computational time is consistently reduced (Figure 3 a–e), up to 24.4%.

## Supplementary

### Replication of the main results with different data resolution

The results of the main text are based on the ico32 resolution (approximately 4 mm inter-vertex distance), which is one of the most used resolutions in surface-based analysis. The other commonly used resolution is ico64 (approximately 2 mm). We repeated our analysis with ico64 resolution data and observed similar results, which we summarize as Supplementary Figures S1–S3, which follow the same structure as Figures 1–3.

**Figure S1.**
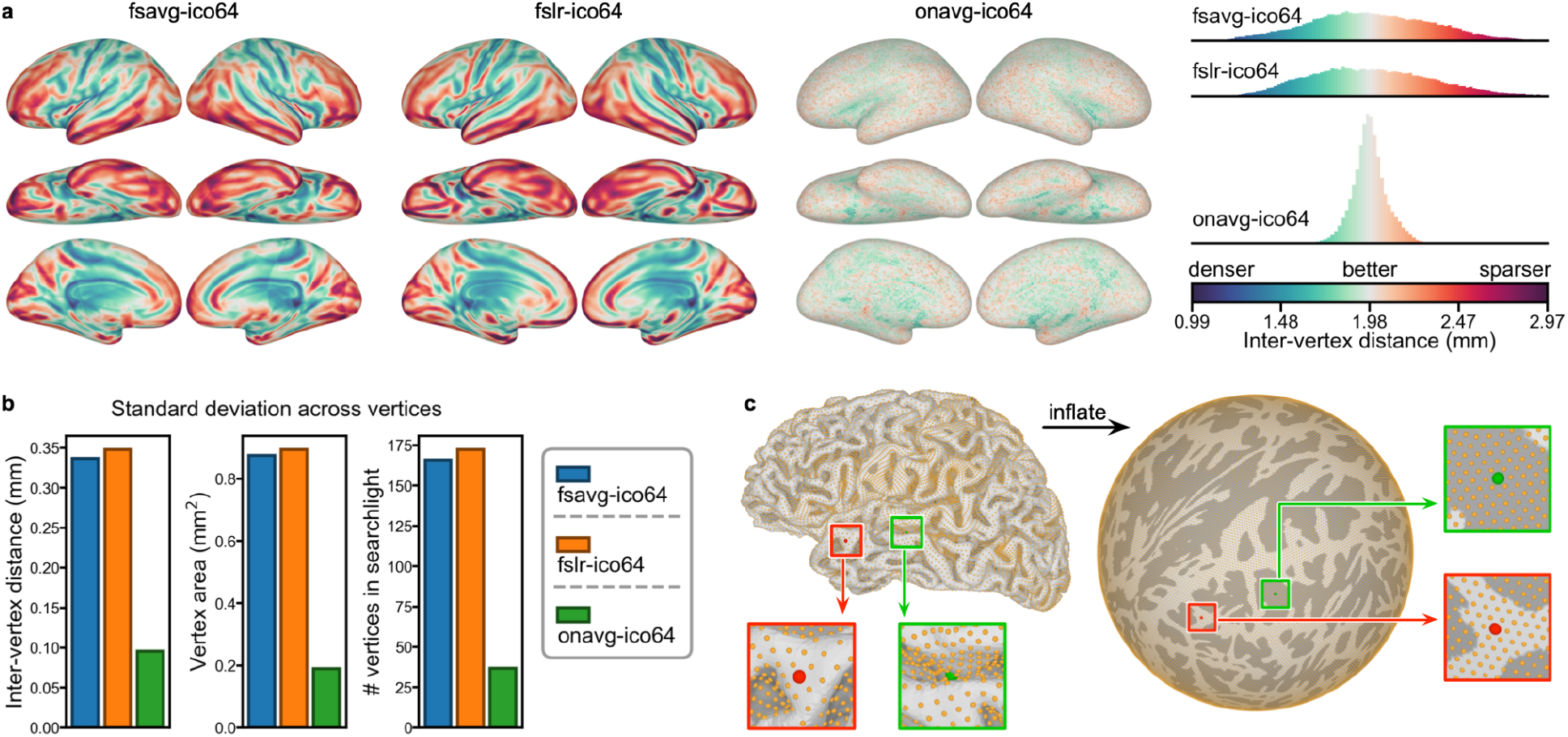
Variation in vertex properties across the cortex at ico64 resolution. For all vertex properties, the variance across the cortex was also much smaller for onavg compared to other templates based on the ico64 resolution (mean = 5.68%; all < 8.04%): For inter-vertex distance, the variance across vertices for onavg was 8.03% compared to fsavg, and 7.52% compared to fslr. For vertex area, the variance for onavg was 4.64% compared to fsavg, and 4.44% compared to fslr. For the number of vertices in a 20 mm searchlight, the variance for onavg was 4.90% compared to fsavg, and 4.54% compared to fslr.

**Figure S2.**
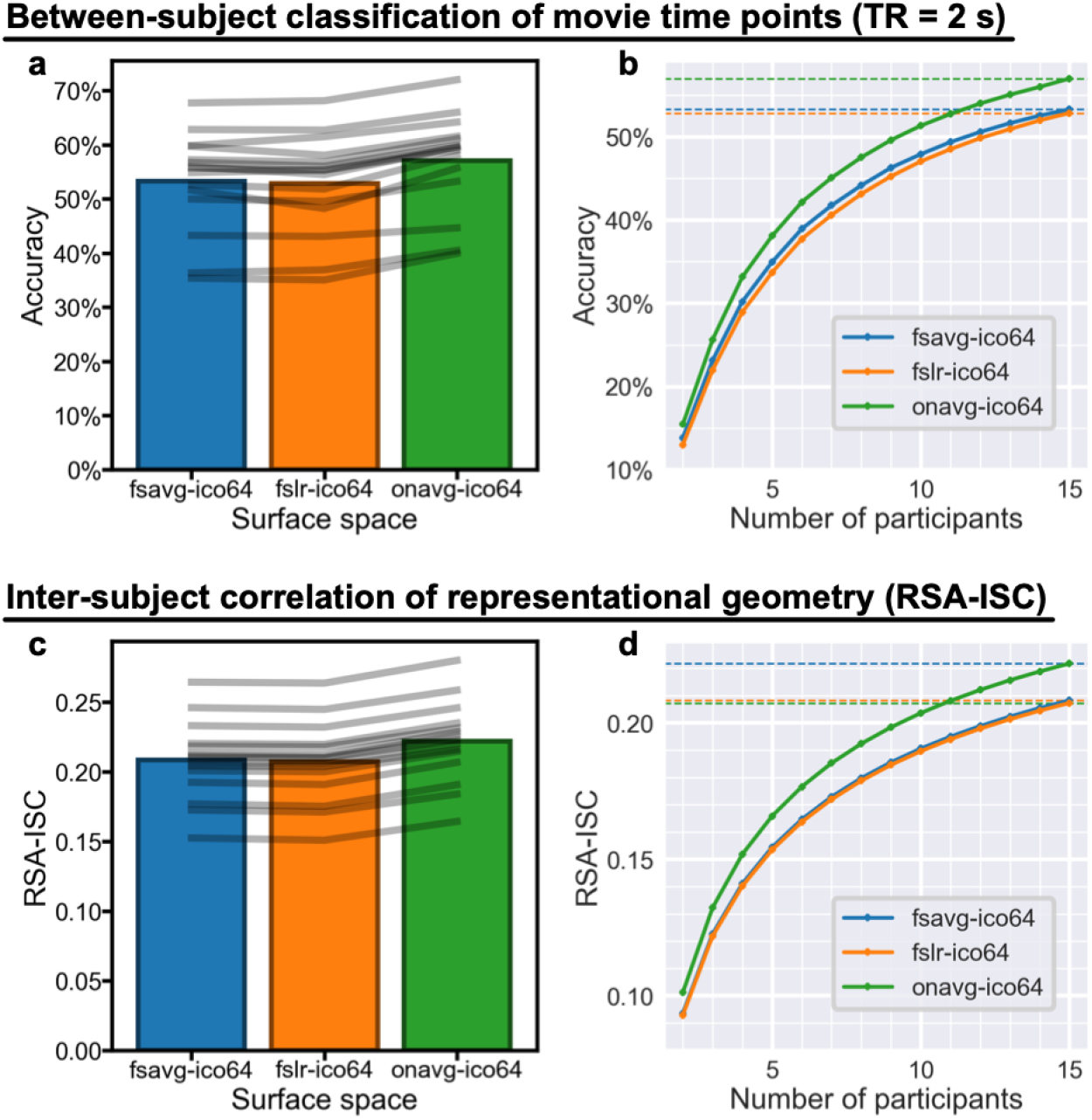
Better cortical sampling improves MVPA results (ico64 resolution). (a) The between-subject classification accuracy of movie time points based on the onavg surface template (57.0% on average) was consistently higher than the accuracy based on fsavg (53.3%) and fslr (52.9%) across all 15 participants. Gray lines denote individual participants. (b) The accuracy based on 15 participants using fsavg or fslr can be achieved by 11–12 participants using the onavg template (52.7% for *n* = 11; 54.0% for *n* = 12). Dashed horizontal lines denote accuracies when *n* = 15. (c) RSA-ISC, computed as the correlation between one participant’s RDM and the average of others’, was also higher when using the onavg template (*r* = 0.222, averaged across searchlights) compared with fsavg (*r* = 0.208) and fslr (*r* = 0.207), and the effect was observed for all 15 individual participants. (d) The noise ceiling based on *n* = 11 and the onavg template (*r* = 0.208) is approximately the same as that based on *n* = 15 and other templates. Across both classification and RSA analyses, we found that the advanced cortical sampling of the onavg template led to more efficient use of fMRI data and better MVPA results, and by switching to onavg the same results can be achieved with less than 3⁄4 of the original sample size.

**Figure S3.**
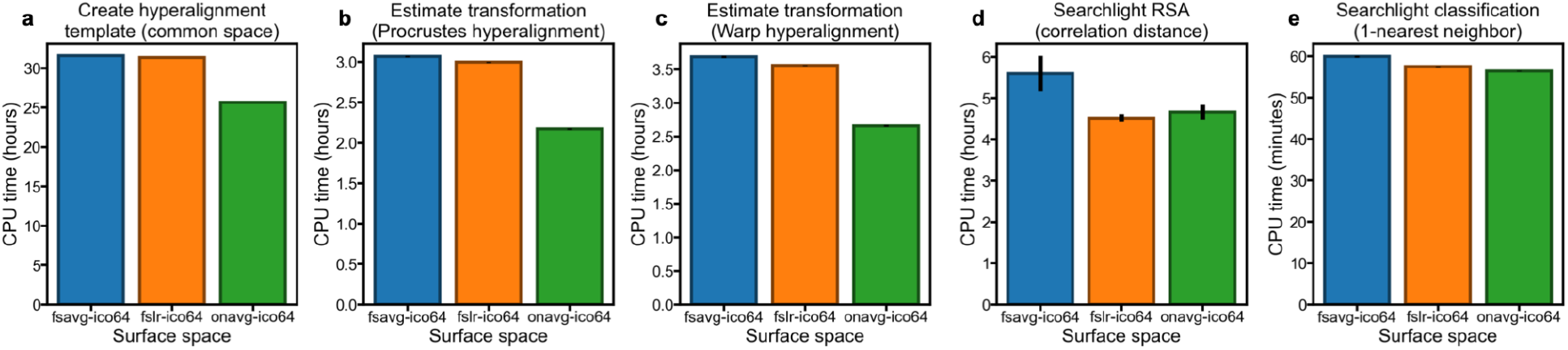
Computational time of representative computational algorithms (ico64 resolution). The anatomy-based uniform sampling of the onavg template avoids sizable searchlights (Figure 1e, Supplementary Figure S1e), resulting in expedited computations for various searchlight-based algorithms, including creating hyperalignment common space (a), aligning individual subjects to the common space (b and c), representational similarity analysis (d), and multivariate pattern classification (e).

### Replication with alternative alignment methods

For each surface template space, we prepared three sets of data based on different alignment methods. The results in the main text are based on warp hyperalignment, and we also repeated the analysis with Procrustes hyperalignment and surface alignment (Supplementary Figures S4 and S5, respectively). We observed a similar advantage of the onavg surface template than other surface templates: Both classification accuracy and RSA-ISC were highest for warp hyperaligned data, as a result of better functional alignment across individuals; the results based on the onavg template were consistently better than other templates for all 15 participants, regardless of the alignment method.

**Figure S4.**
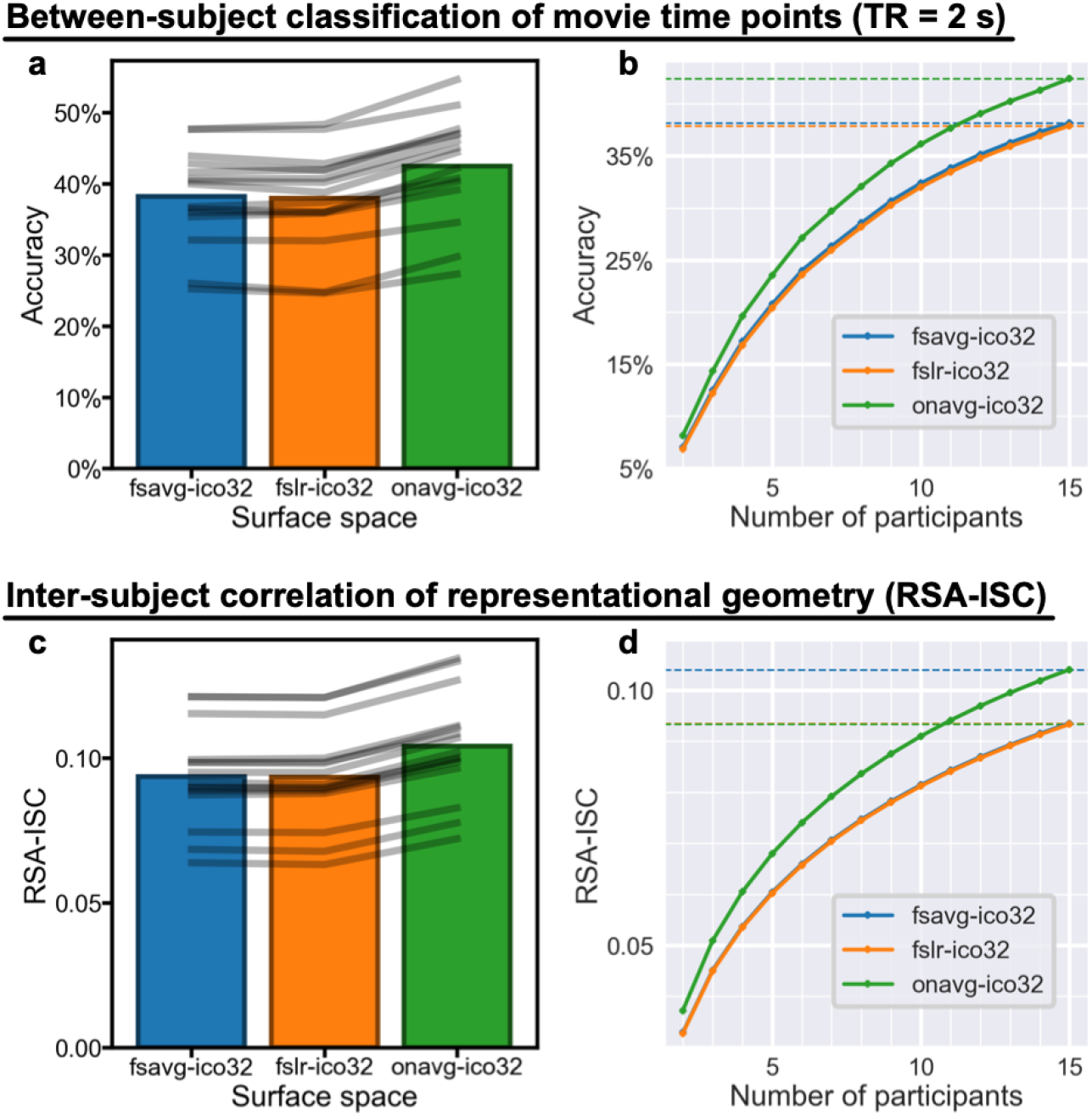
Better cortical sampling improves MVPA results (Procrustes hyperalignment). (a) The between-subject classification accuracy of movie time points based on the onavg surface template (42.4% on average) was consistently higher than the accuracy based on fsavg (38.1%) and fslr (37.9%) across all 15 participants. Gray lines denote individual participants. (b) The accuracy based on 15 participants using fsavg or fslr can be achieved by 11–12 participants using the onavg template (37.7% for *n* = 11; 39.0% for *n* = 12). Dashed horizontal lines denote accuracies when *n* = 15. (c) RSA-ISC, computed as the correlation between one participant’s RDM and the average of others’, was also higher when using the onavg template (*r* = 0.104, averaged across searchlights) compared with fsavg (*r* = 0.094) and fslr (*r* = 0.093), and the effect was observed for all 15 individual participants. (d) The noise ceiling based on *n* = 11 and the onavg template (*r* = 0.094) is approximately the same as that based on *n* = 15 and other templates. Across both classification and RSA analyses, we found that the advanced cortical sampling of the onavg template led to more efficient use of fMRI data and better MVPA results, and by switching to onavg the same results can be achieved with less than 3⁄4 of the original sample size.

**Figure S5.**
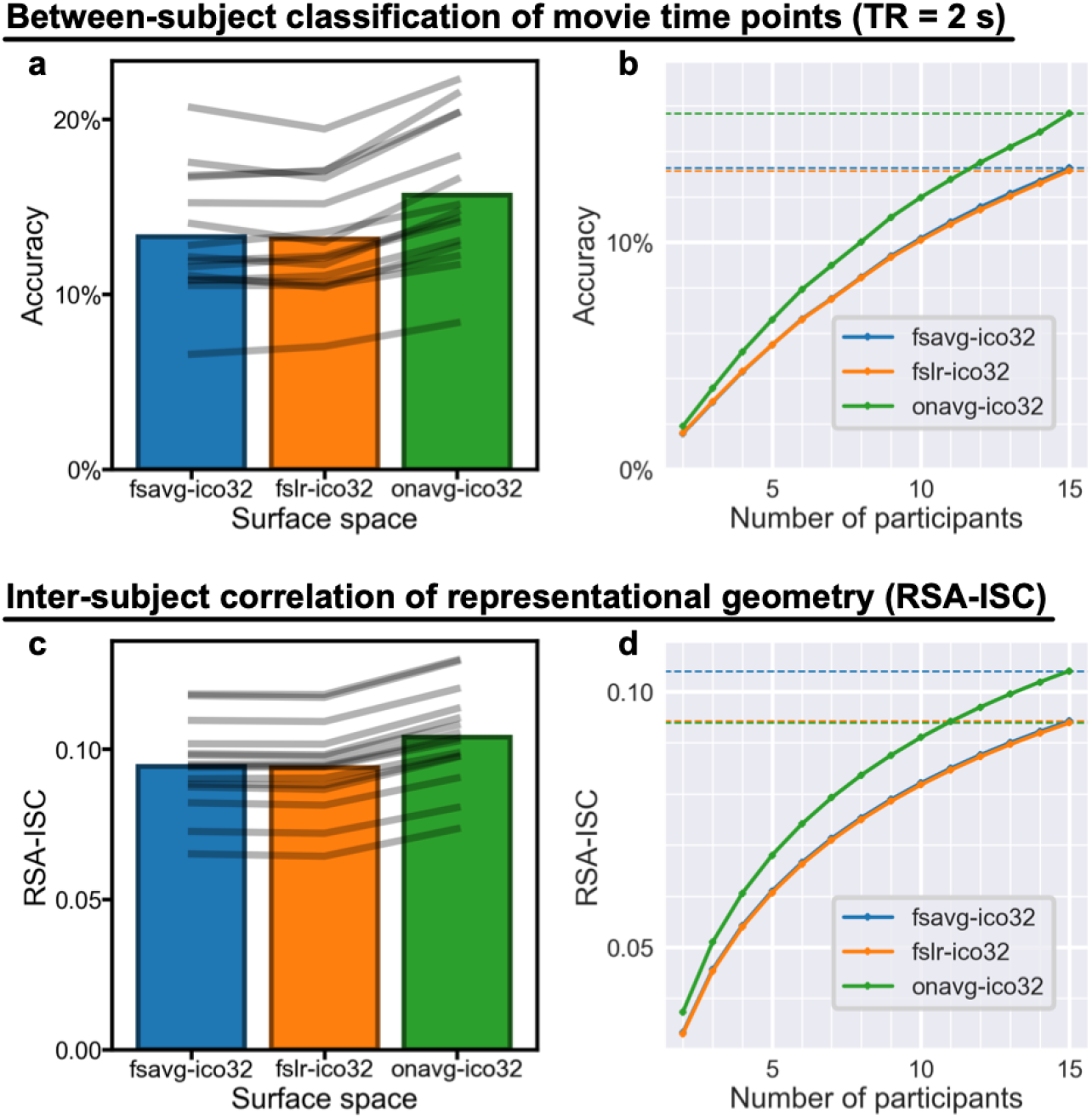
Better cortical sampling improves MVPA results (surface alignment). (a) The between-subject classification accuracy of movie time points based on the onavg surface template (15.7% on average) was consistently higher than the accuracy based on fsavg (13.3%) and fslr (13.2%) across all 15 participants. Gray lines denote individual participants. (b) The accuracy based on 15 participants using fsavg or fslr can be achieved by 11–12 participants using the onavg template (12.8% for *n* = 11; 13.5% for *n* = 12). Dashed horizontal lines denote accuracies when *n* = 15. (c) RSA-ISC, computed as the correlation between one participant’s RDM and the average of others’, was also higher when using the onavg template (*r* = 0.104, averaged across searchlights) compared with fsavg (*r* = 0.094) and fslr (*r* = 0.094), and the effect was observed for all 15 individual participants. (d) The noise ceiling based on *n* = 11 and the onavg template (*r* = 0.094) is approximately the same as that based on *n* = 15 and other templates. Across both classification and RSA analyses, we found that the advanced cortical sampling of the onavg template led to more efficient use of fMRI data and better MVPA results, and by switching to onavg the same results can be achieved with less than 3⁄4 of the original sample size.

### Replication with different amounts of data

We repeated our analysis with smaller numbers of participants, and we found a consistent advantage of onavg over other surface templates, regardless of the number of participants. In Supplementary Figures S6 and S7, the curves indicate the number of participants for onavg (*y*-axis) and other templates (*x*-axis) to achieve the same classification accuracy and RSA-ISC. The solid black lines indicate *y* = *x*, and the dashed black lines indicate *y* = ¾ *x*. The curves are all close to the dashed line in all 12 panels, which indicates onavg only needs 3/4 of the amount of data to achieve the same performance as other surface templates for MVPA algorithms, different alignment methods, different data resolutions, and different numbers of participants.

**Figure S6.**
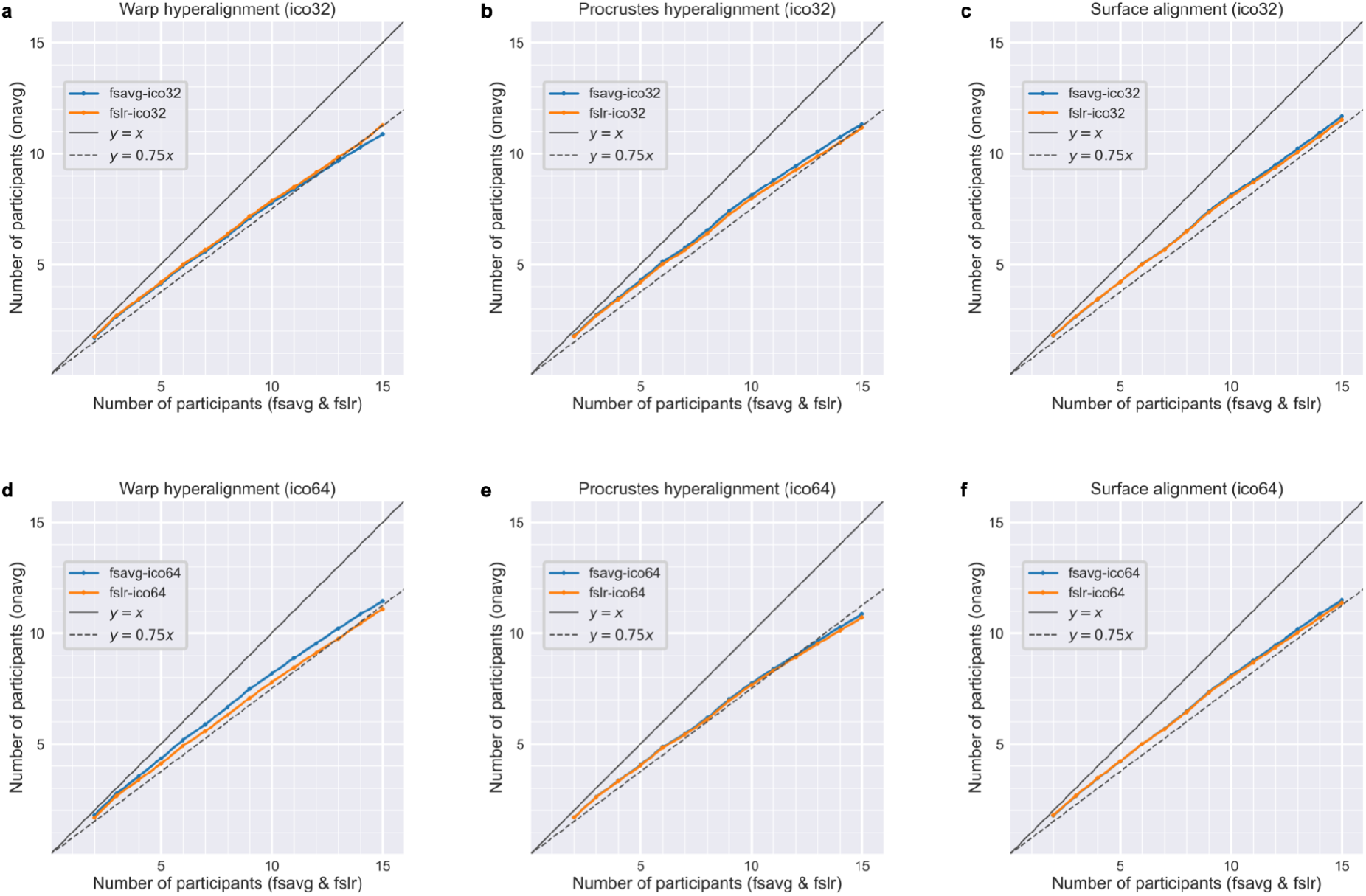
The number of participants for onavg (*y*-axis) and other templates (*x*-axis) to achieve the same classification accuracy. Panels in different rows are based on different data resolutions (top: ico32; bottom: ico64), and panels in different columns are based on different alignment methods (left: warp hyperalignment; middle: Procrustes hyperalignment; right: surface alignment). Across all six panels, there is a consistent pattern that onavg only needs approximately 3/4 of the amount of data to achieve the same multivariate pattern classification accuracy (dashed black line).

**Figure S6.**
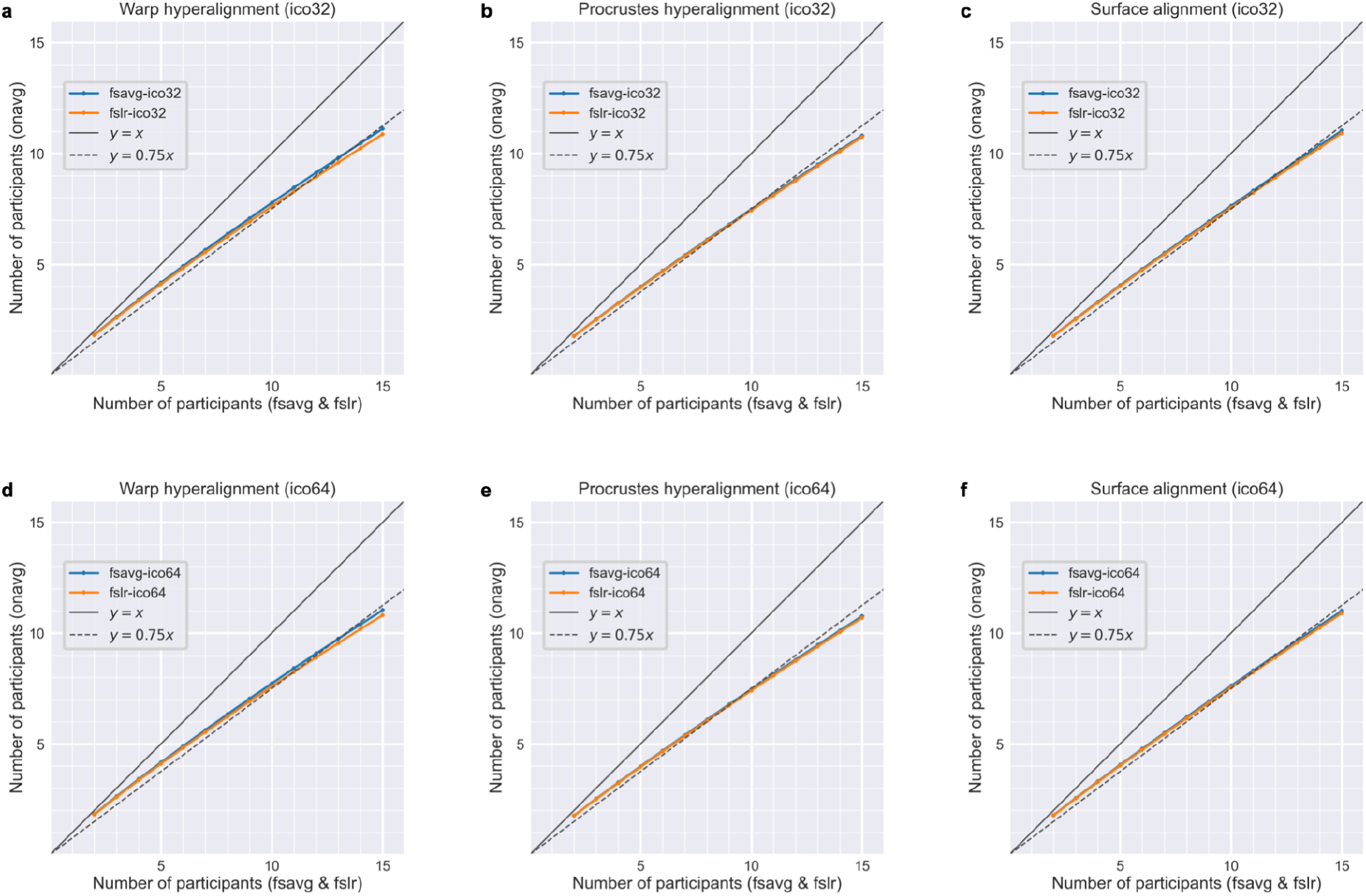
The number of participants for onavg (*y*-axis) and other templates (*x*-axis) to achieve the same RSA-ISC. Panels in different rows are based on different data resolutions (top: ico32; bottom: ico64), and panels in different columns are based on different alignment methods (left: warp hyperalignment; middle: Procrustes hyperalignment; right: surface alignment). Across all six panels, there is a consistent pattern that onavg only needs approximately 3/4 of the amount of data to achieve the same inter-subject correlation of RDMs (dashed black line).

### Improvement by brain region

**Figure S8.**
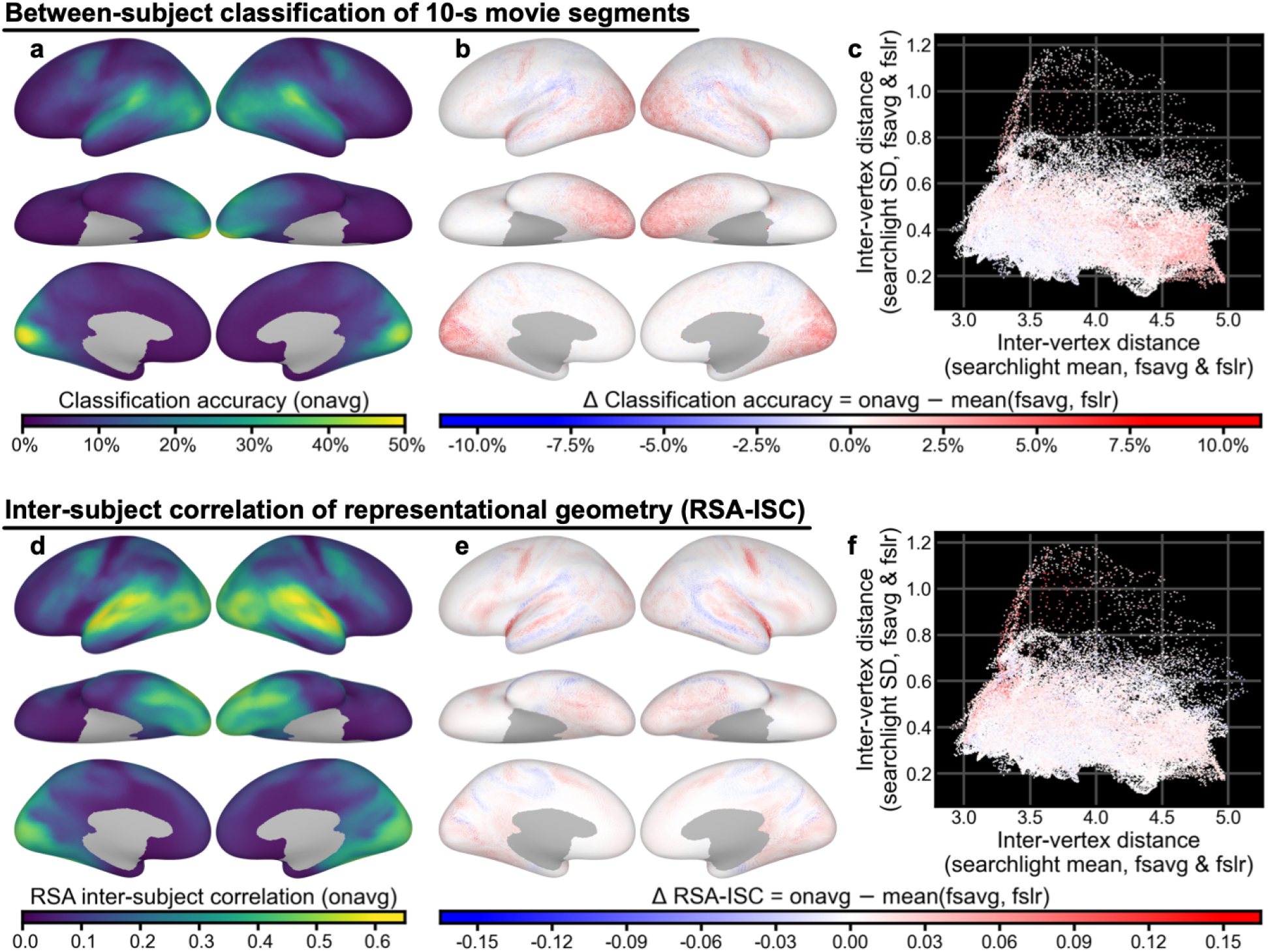
Improvement by brain region. To locate the brain regions that benefit most from anatomy-based sampling and the onavg template, we performed the classification analysis and RSA analysis searchlight by searchlight for each surface template and compared the results across surface templates. Consistent with our previous results, the occipital, lateral temporal, ventral temporal, and lateral prefrontal regions had the largest classification accuracy (a) and RSA-ISC (d) among all brain regions. Classification accuracy (b) and RSA-ISC (e) for much of the cortex were higher based on onavg than on other surface templates. For the classification analysis, the improvement was most prominent in regions that were sparsely sampled by other templates (c), such as the occipital cortex and ventral temporal cortex. For the RSA analysis, the improvement was most prominent in regions that were inhomogeneously sampled by other templates (f), such as the lateral temporal cortex and the lateral prefrontal cortex.

**Figure S9.**
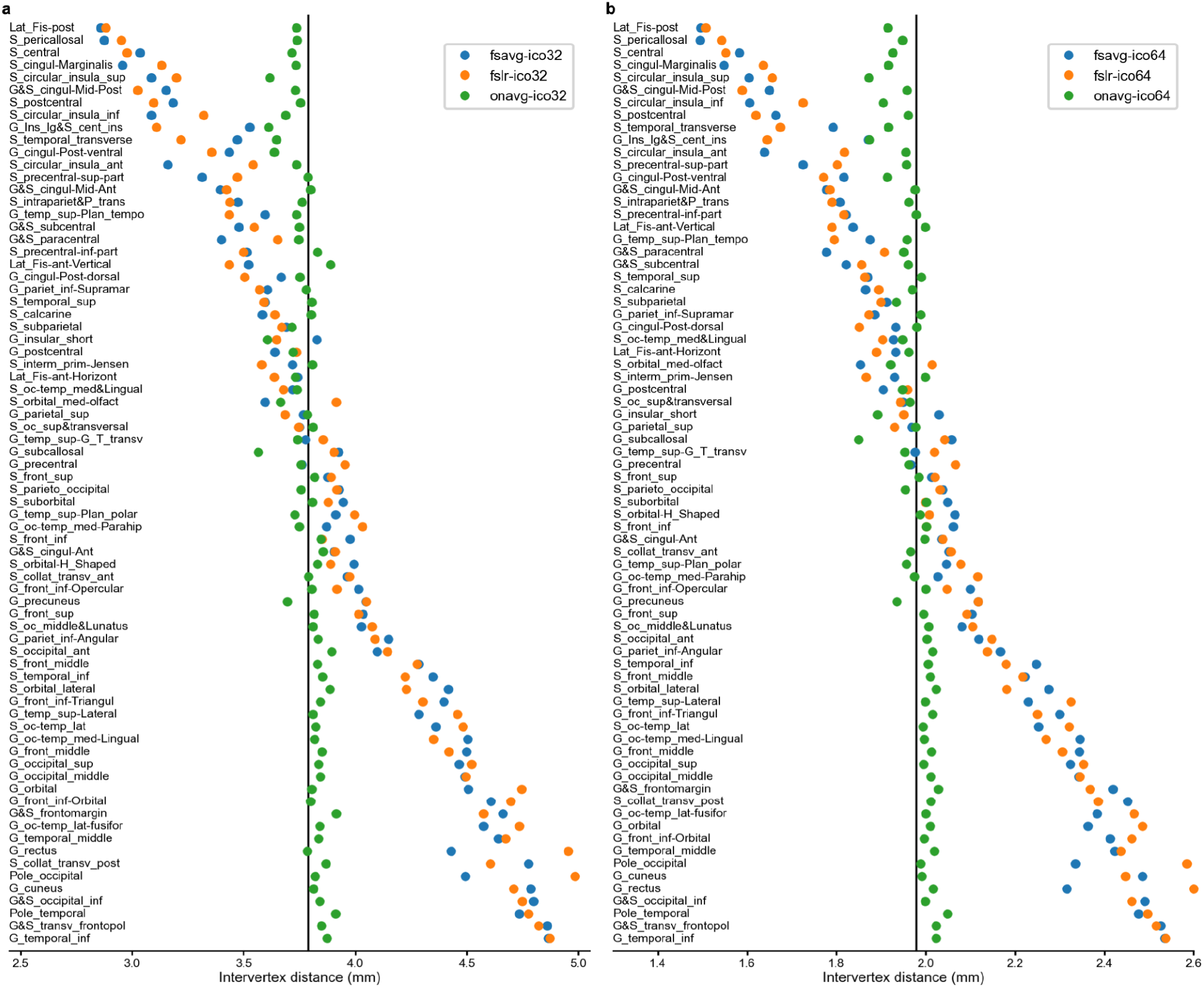
Inter-vertex distance by brain region. Traditional surface templates sample the cortex based on the spherical surface, which results in systematically differential sampling rates across regions due to geometric distortions, measured by inter-vertex distance. For example, the posterior part of the lateral fissure (Lat_Fis-post) is densely sampled by both fsavg and fslr for both ico32 and ico64 resolutions; in contrast, the inferior temporal gyrus (G_temporal_inf) is sparsely sampled across these conditions. The issue of uneven sampling is minimized by the onavg template space (green dots), whose vertices were chosen based on the anatomical surface instead of the spherical surface.

## Notes

https://github.com/feilong/tpl-onavg

### Competing Interest Statement

The authors have declared no competing interest.

